# Seasonal migration as a life history trait facilitating adaptation to climate change

**DOI:** 10.1101/2021.09.01.458412

**Authors:** Katherine Carbeck, Tongli Wang, Jane Reid, Peter Arcese

## Abstract

Predicting the geographic range of species and their response to variation in climate are entwined goals in conservation and evolutionary ecology. Species distribution models (SDMs) are foundational in this effort and used to visualize the geographic range of species as the spatial representation of its realized niche, or when based only on climate, its climate niche. SDMs can also forecast shifts in species range given climate change, but often lack of empirical support for causal links between climate and demography, yielding uncertain predictions. We addressed such uncertainties whilst also exploring the role of migration and resident life-histories in climate adaptation in mobile animals using 48 years of detailed demographic and climate data for song sparrows (*Melospiza melodia*), a polytypic species that varies in migratory life history. We developed SDMs representing demographic and climate niches of migratory and resident populations in western North America from California (CA) to Alaska (AK) using data from a focal population in British Columbia (BC) and 1.2 million citizen science observations. Distributions of resident and migrant populations predicted by each model agreed strongly (72.8%) in the region of our focal population, but less well in regions with dissimilar climates. Mismatches were largest in CA, smaller in AK, but in all cases supported the hypothesis that climate influences the evolution of migration and limits year-round residency. Our results imply that migrants predominated in our focal population a century ago, but that climate change has favored range expansions by non-migratory phenotypes and facilitated an upward shift in the elevational range of residents. We suggest long-term studies are crucial to evaluating the predictions of SDMs positing causal links between climatic conditions and species demography. We found such links to be robust regionally and particularly useful to elucidating the potential for migration or residence to facilitate adaptation to climate change.

## Introduction

Theory and empirical evidence indicate that spatial and temporal heterogeneity in the environment can drive variation in species distribution and life history by causing variation in individual fitness and thereby driving natural selection and adaptation to local conditions (Wright, 1982; Wade & Kalisz, 1990; Schluter, 2000; Aitken et al., 2008; Hendry et al., 2018). However, the rapid pace of anthropogenic change has raised new questions about the capacity of local populations to accommodate change (e.g., Gaston 2003; Mason et al., 2019; reviewed by Root et al., 2003; Parmesan, 2006; Bell, 2017) and elevated the importance of empirical studies which allow us to estimate the pace of climate adaptation and identify the processes underlying it (e.g., Grant & Grant, 2006; Stoks et al., 2014; Bay et al., 2018; Bontrager & Angert, 2019; Radchuk et al., 2019). Theory suggests that species can avoid the negative effects of climate change on demographic performance at a site by (1) adapting via genetic mechanisms that increase individual fitness and facilitate persistence at sites occupied historically (Aitken et al., 2008; Bay et al., 2018); (2) adapting to current conditions via phenotypic plasticity (i.e. phenotypic change within existing genotypes, Ghalambor et al., 2007; Charmantier et al., 2008; Nicotra et al., 2010); and/or (3) dispersing to new areas that now match historical climatic conditions (Angert et al. 2011; Chen et al. 2011; Gillings et al. 2015; *cf* Greenwood & Harvey 1982). However, many species also harbor capacities to respond to deteriorating environments by altering spatial patterns of seasonal migration (i.e., the reversible movement between breeding and non-breeding areas) versus year-round residency; changes which may arise via adaptive migratory plasticity and/or evolution (Reid et al. 2018). Here, we develop and test the hypothesis that migration represents an additional mechanism by which mobile species can adapt to climate change by facilitating their access to spatially and temporally limiting resources whilst avoiding conditions that would preclude their survival at one location throughout the annual cycle.

Populations comprising a mix of seasonal migrants and year-round residents occur in diverse taxa (e.g., fish, Chapman et al., 2012; birds, Dingle, 2014; mammals, Avgar et al., 2014; and reptiles and amphibians, Shaw & Levin, 2011; Yackulic et al., 2017) and may harbor sufficient genetic variation and/or plasticity in migration to enable rapid responses to environmental change (Liedvogel et al., 2011). Such responses can be driven by spatial and temporal variation in the costs and benefits of seasonal migration, which may in turn give rise to correlations between environmental conditions, occurrence of migration, and life history evolution. For example, climate warming and supplemental feeding have each facilitated the establishment of resident populations in species of birds known previously as obligate migrants (Zuckerberg et al., 2011; Møller et al., 2014; Shephard et al., 2015; Plummer et al., 2015). Such transitions can elevate population growth rate in newly-established resident as compared to migrant sub-populations by enhancing over-winter survival and annual reproductive rate (Visty et al., 2018), and similar trade-offs involving individual fitness and climate can drive variation in migratory phenotype across populations. For example, Acker et al. (2021a; 2021b) reported context-dependent fitness of residents and migrants in European shags (*Phalacrocorax aristotelis*), wherein selection favored migrants in severe winters but residents in milder years.

In the context of climate change, such observations suggest that we should also observe temporal shifts away from the expression of seasonal migration and in favor of year-round residence in populations where climate change has ameliorated previous limits on population growth. Similarly, shifts from residence to migration might be expected in areas occupied historically by residents, but which have become inhospitable due to the negative effects of climate change on habitat, food supply, or interacting species. However, documenting such changes requires detailed knowledge of temporal and spatial variation in the occurrence of migration in populations and the demography and growth rate of such populations at large geographic ranges; two requirements rarely met in free-living species. Alternatively, one can infer continent-wide variation in the growth rates of resident versus migrant populations by extrapolating from long-term climate and demographic data collected at smaller scales. However, because datasets spanning multiple decades and completely enumerated populations are rare, it remains to be demonstrated that temporal variation in climatic factors shown to limit population growth at local scales can be reliably scaled-up to predict the distribution of resident and migratory populations of a species across its historical, contemporary and/or future range. Using such relationships can predict species distribution and evaluating the reliability of those predictions at regional and larger scales thus remain key steps to elucidating mechanisms by which mobile species may respond change in the environment.

Most efforts to predict the response of species to environmental change employ species distribution models (SDMs) to summarise statistical links between the occurrence of species and environmental characteristics at a site (reviewed by Elith & Leathwick, 2009). Climate niche models are a particular case of SDMs that aim to predict species’ distribution based solely on climatic factors known or assumed to limit population growth (Andrewartha & Birch, 1954; Wiens et al., 2009). However, such models have been criticized for their opaque assumptions and inability to reveal the biological mechanisms driving variation in demographic performance and species distribution (Elith & Leathwick, 2009; Zurell, 2017). For example, although climate niche models make implicit predictions about population growth in versus outside the climate niche, few studies test such assumptions directly (e.g., Tredennick et al., 2017; Bayly & Angert, 2019; Williams et al., 2021). Moreover, eco-evolutionary studies increasingly indicate that local adaptation to environment is common in nature (e.g., Hoban et al. 2016; Delph 2018), implying that some species can be expected to show heterogeneous responses to climatic factors at micro-geographic to regional scales (e.g., Walsh et al., 2019; Mikles et al., 2020). Because many climate niche models assume that the biological and statistical relationships between climate and demography act similarly over the species range, testing their underlying mechanisms and predictions will be essential to evaluating their reliability, evaluating the adaptive capacity of species, and predicting how resident and migrant populations may respond to spatial and temporal heterogeneity in climatic conditions.

We addressed the knowledge gaps above using 48 years of demographic data, 1.2 million citizen science observations, and climate data spanning two centuries to predict the influence of historical, contemporary, and future climate on population growth, and migration versus residence, in song sparrow populations throughout western North America (comprising Alaska, British Columbia, Washington, Oregon, and California). To do so, we first tested several *a priori* hypotheses (Table 1) on the effects of temporal variation in climate on survival and reproduction in a focal study population which resides year-round in coastal British Columbia, Canada.

**Table 1.**
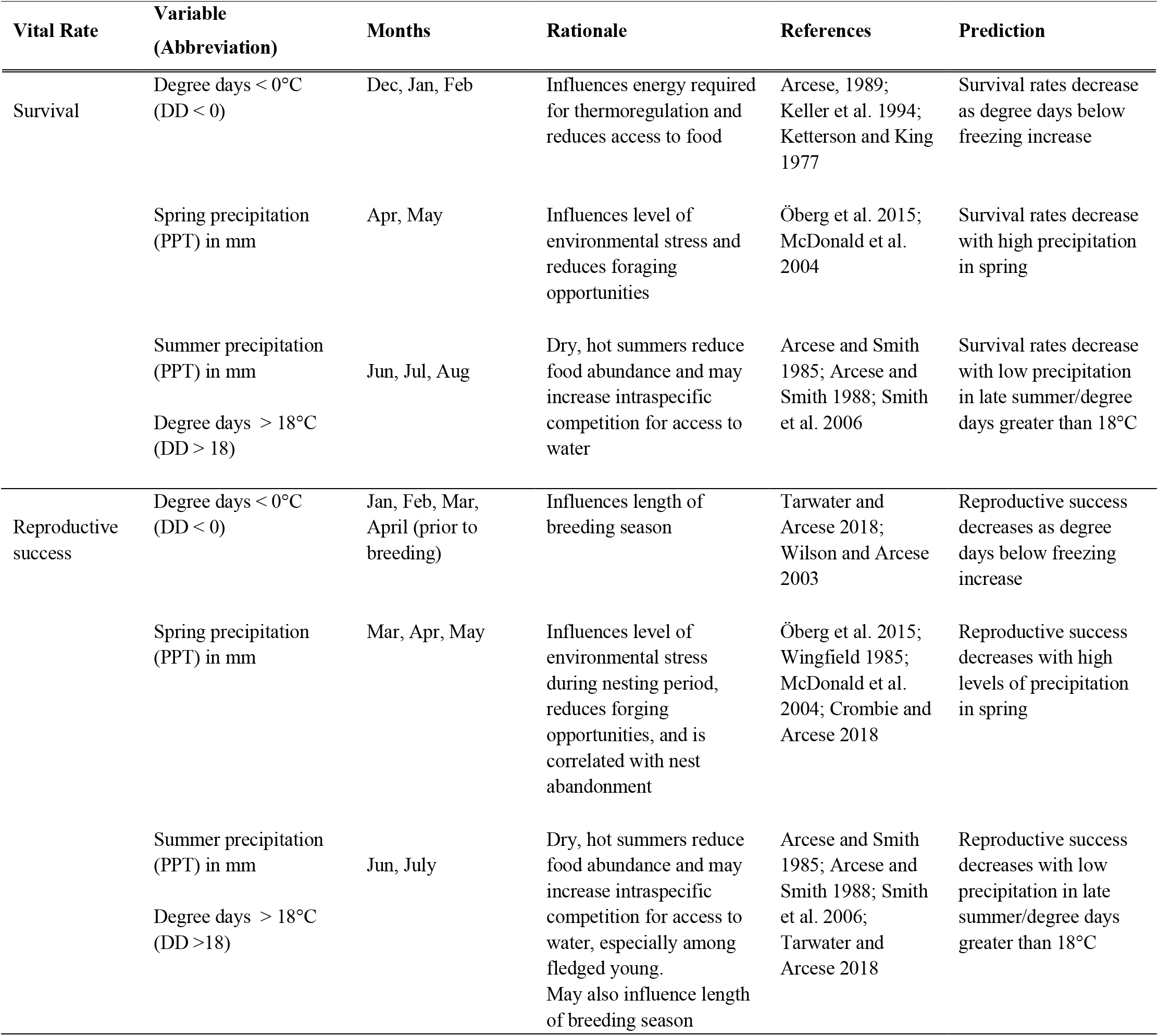
Predicted response of demographic vital rates to climatic variables, their hypothesized mode of action, and references to the prior results on which predictions are based.

Second, we then used ‘space-for-time substitution’ (e.g., Pickett 1989; La Sorte et al. 2009) to predict how variation in climate may affect the population growth rate (λ), migratory status, and distribution of song sparrow populations (i.e., the ‘demographic niche’) at larger spatial scales, assuming that song sparrows in our study population and western North America respond similarly to climate. This study area encompasses Mediterranean to Arctic ecosystems, an elevational range of 0 to 3200 m, and 17 of 25 extant subspecies (Miller, 1956; Patten & Pruett, 2009), including five of ‘special concern’ (Pruett et al., 2008; Shuford et al., 2008). Because climate is thought to play a key role in local adaptation and the evolution of migration or residence in song sparrows (e.g., Aldrich 1984, Arcese et al. 2002, Mikles et al. 2020), they are an excellent species in which to develop and test SDMs.

Third, we evaluated the performance of our demographic niche model by comparing it to a climate niche model generated from 1.2 million citizen science observations and contemporary climate data to predict the distributions of migratory and resident populations throughout western North America. Given the large scale of our study area and substantial variation in climate recorded therein, we expected that mismatches between our demographic and climate niche models would increase as climatic conditions diverged from those experienced in our focal study population and reveal practical limits to prediction.

Last, we predicted how climate might influence migratory phenotype by using our demographic niche model to predict the historical and future ranges of migratory and resident song sparrow populations in three periods over two centuries: 1901-1910, when winter temperatures in our focal study population averaged 0.9°C cooler than present-day (2010-2018), and 2070-2100, when winter temperatures are predicted to exceed conditions 200 years earlier by 3.1°C. We validated retrospective projections using historical records before speculating on the potential role of migration versus residence in climate adaptation. By doing so, we provide a particularly detailed case study of how population demography, species occurrence, and climate data can be used to explore, predict, and test for the interactive effects of environmental change on the distribution, demography, and migratory dynamics of mobile species.

## Methods

### Study Species

Twenty-four subspecies of song sparrows breed from Newfoundland to the Aleutian Islands, and south to central Mexico, with 17 subspecies occurring with our study area from Alaska (AK) to California (CA). Variation in size and plumage over the song sparrow range exemplify Bergmann’s and Gloger’s rules, with the largest individuals residing year-round in near-shore habitats of the Aleutian Islands and the smallest in California salt marshes, and the darkest inhabiting southeast Alaska and lightest the arid southwest of North America (Aldrich, 1984; Zink & Remsen, 1986; Patten and Pruett, 2009; Pruett & Winker, 2010). Because song sparrows are among the world’s most polytypic species, display marked local adaptation, and additive genetic variation in traits linked to environmental conditions (e.g., Walsh et al., 2019; Mikles et al., 2020), they are an excellent species to test for spatial and temporal heterogeneity in relationships between climate, population growth, and migration.

### Focal population

All data on the demography of song sparrows was collected on Mandarte Is., BC, Canada (48.8°N, 123.8°W; *c*. 6 ha in area), where all song sparrows were identified individually and monitored from 1960-1962 and 1975-2018 using nearly identical methods (Tompa, 1963; Smith et al., 2006). Briefly, territories were monitored each 2-5 days from mid-March to July in all years to record the behavior, survival, and reproductive success of all birds. Nestlings were uniquely color-banded after hatching, observed to independence from parental care (∼24-32 days of age), and recorded as having recruited to or disappeared from the population in late April the following year. High annual re-sighting probabilities (> 99%; Wilson et al., 2007), the enumeration of immigrants by color-banding (< 0.5 female/yr on average; Reid & Arcese 2020), continuous monitoring of breeding activity, and genetic confirmation of a 50:50 sex ratio at hatching (Postma et al., 2011) facilitated high precision in our estimates of survival, reproduction, and population growth. For simplicity, we only considered females when estimating demographic rates and population growth here (Arcese et al., 1992; Arcese & Marr, 2006). Smith et al. (2006), Sardell et al. (2012). Lameris et al. (2016) describe the current and historical change in vegetation on the island.

### Climate data

All climate data were obtained via ClimateNA (version 6.00; Wang et al. 2016), which generates scale-free point data for specific locations through dynamic local downscaling of gridded historical and future climate variables for individual years and periods between 1901 and 2100. Historical and contemporary periods were selected based on the most recently available decadal data from ClimateNA. For building climate niche and demographic models, climate data were generated for specific sample locations for the contemporary period (2010-2018). For spatial predictions, monthly climate data obtained by querying ClimateNA using an input file of the latitude, longitude and elevation each of 35,118,256 rasterized cells in our study area (1 km^2^ DEM; Amatulli et al., 2018). Historical (1901-1910) and future climate (2070-2100) was estimated similarly. To account for uncertainty in future climate, we selected an ensemble of 15 General Circulation Models (GCM; Coupled Model Intercomparison Project) included in the IPCC Fifth Assessment Report (IPCC, 2014), and an intermediate greenhouse gas emission scenario (RCP 4.5) which assumes emissions start declining around 2040 (IPCC, 2014).

### Demographic niche models

We characterized the effects of monthly weather on demographic performance using variables shown previously to predict adult survival (S_a_), juvenile survival (S_j_), reproductive success (RS), and population growth rate (λ) (Table 1, S1), and thought to reflect extreme winter and/or summer conditions influencing energy and water balance, respectively (e.g., Arcese et al., 1992; Wilson & Arcese, 2003; Tarwater & Arcese, 2018). S_a_ was estimated annually as the fraction of females alive in late April each year that survived to the next April. S_j_ was estimated as the fraction of female yearlings that became independent from parental care and survived on Mandarte to late April the next year. RS equaled the mean number of female young that became independent of parental care per adult female in each year (see Fig. S1 for timeline). Population growth rate was then calculated as λ = (S_j_ * RS) + S_a_ following Arcese & Marr (2006; see also Visty et al., 2018), assuming no further age structure or immigration.

We employed normal regression to quantify relationships between S_j_, S_a_, RS and climate. All models initially included linear and second-order polynomial terms of each *a priori* predictor identified in Table 1. We then reduced models using supervised backward selection to eliminate variables sequentially with the least influence on model fit, such that final models only included predictors with influence (p < 0.1) that were not closely correlated (r < 0.7). All response variables were mean-centered and natural log-transformed to facilitate climate mapping and because doing so led us to models that explained more variance and exhibited better diagnostics than other approaches (Table S3). Values of S_j_ and S_a_ predicted from generalized linear models assuming binomial errors and a logit link were well correlated to values derived above (*r* = 0.99 and 0.770, respectively); we employed the former here because similar methods generated our *a priori* predictions (Table 1).

We then used our fitted demographic models to predict contemporary λ across our study area given climate (Table S3; see ‘Climate data’). To do so, predicted values of S_a_, S_j_, and RS values were back-transformed, centered on their observed means in our focal population, and bounded between 0 and 0.99 for S_a_, 0 and 0.69 for S_j_, and 0 and 4.34 for RS, which represent ranges spanning ± 3 SEs. We used the same procedure to predict historical and future λ in relation to climate. Each map cell was classified as supporting a ‘resident’ population if the predicted value of λ predicted given local climate at the site was ≥ 1 (i.e., resident demographic niche), or a ‘migrant’ population if λ < 1 (i.e., migrant demographic niche), given our assumption that the long-term persistence such populations at a site can only be achieved via seasonal migration.

### Climate niche models

To characterize relationships between migrant or residence and climate using our ‘climate niche model,’ we first obtained all observations of song sparrows from the eBird Basic Dataset (version Sep 2019; www.ebird.org/science/download-ebird-data-products) to extract presence and absence data from individual checklists. eBird is a large semi-structured citizen science depository of high-quality observations of birds year-round. We filtered 49,972,482 checklists for our study area (comprising Alaska, British Columbia, Washington, Oregon, and California) to include only those recorded from January 1, 2010 to September 1, 2019, representing ‘complete checklists’, with a maximum distance travelled of 5 km, and ≤ 5 hrs of effort, and complete documentation; yielding 5,137,845 informative checklists (using *Auk* package in R; Strimas-Mackey et al., 2018). Because eBird checklists are not randomly distributed in space or time and often suffer ‘class imbalance’ due to more absences than presences recorded, we generated a 1 km hexagonal grid over the study area and randomly subsampled the filtered dataset to reduce bias, following Johnston et al. (2019). Subsampling had little influence on prevalence but reduced checklists to 3,424,036 (detection rate before subsampling = 33.3%; after= 33.9%). Checklists were further sorted to create ‘winter’ and ‘breeding’ distributions based on the date in which observations were made, such that observations in January-February were assumed to reflect the distribution of overwintering populations (N = 543,837) and those in May-June the distribution of the breeding populations (N = 693,751), respectively (N_total_ = 1,237,588). Ten annual climate variables related to temperature and precipitation were generated for each of the sample locations. Those climate variables were expected to influence the occurrence of migration and residence in song sparrows (Table S2; 2010-18).

We adopted Random Forest (*ranger* R package; Wright & Ziegler, 2017) to predict the contemporary, historical, and future ranges of migratory and resident song sparrow populations in western North America. The species occurrence (presence or absence) was used as the dependent variable, and the 10 climate variables were used as predictors. Random Forest works by producing a ‘forest’ of decision trees and aggregating the results over all trees. The decision trees are constructed with a bootstrap sample of the input data such that the resulting ‘bagged’ dataset contains about 64% of the original observations, and the remaining samples comprise the ‘out-of-bag’ (OOB) data. Using the trees grown from a bootstrap sample, each of the independent observations in the OOB data is classified as either presence or absence and a model prediction error (OOB error) as the percent of incorrectly classed observations is calculated. Random Forest models are considered one of the most credible statistical methods for species distribution modelling, and are highly flexible and easy to implement (Elith et al. 2008; Iverson et al. 2011; Laube et al. 2015; Wang et al. 2012).

We evaluated OOB error rates using Brier Score (Brier, 1950), which employs the mean squared error of the probabilistic model predictions and the true presence or absence in OOB data. We also evaluated performance using sensitivity, specificity, AUC, and Kappa metrics calculated by comparing predictions to the actual observations in unseen data (OOB data; Table S4). Random Forest also provides measures of variable importance defined as the mean decrease in model accuracy (DMA; Altmann et al., 2010). To reduce prediction bias, we fit a balanced random forest model using the ‘sample.fraction’ argument to grow each tree from random samples of the data with an equal number of detections and non-detections. Because the effect of number of predictors selected at each node was minor, we used the default square-root of the number of predictors included (Breiman, 2001). By repeated testing, we found that 500 trees per model generated consistency in model accuracy.

We first predicted the climate niche for resident and breeding song sparrows using the predict function in the *ranger* package to produce maps in response to the contemporary, historical, and future climate periods (see ‘Climate data’). We then created the migrant niche by classifying ‘migrant’ pixels as those which were occupied during the breeding period and absent in winter. The resident niche was determined by classifying ‘resident’ pixels as those which were occupied during both breeding and winter periods, assuming the same individuals are present in both seasons. All niche models were then visualized in ArcGIS (v10.7.1).

### Spatial and temporal map comparisons

To assess agreement of our climate and demographic niche models we first calculated the percentage of grid cells predicted to host resident or migrant song sparrow populations given the climate niches of each. We then compared these climate niches to the demographic niche model derived from the demographic variation observed on Mandarte Is. and projected across the study area (i.e., assuming that all song sparrows respond to climatic conditions in ways similar to those documented on Mandarte Is. over the last half century). To do so, we also classified grid cells by the expected rate of population growth given local climatic variation as noted above. Agreement between mapped predictions was estimated by first sampling 1,000 grid cells at random and calculating the Pearson’s correlation coefficient, *r*, between the difference in predicted probabilities of occurrence and population growth rate at a site versus the Euclidian distance from our focal location; we report the mean value of *r* (±1SD) given 10,000 replicated comparisons and the mean percentage of grid cells in agreement in three regions of interest at the northern (AK), central (BC), and southern (CA) extents of our study area. Statistical description of seasonal and spatial climate variation in these regions appear in Fig S5 and Table S6.

We also estimated the correlation between maps in climatic space by collapsing the 10 climate variables (Table S2) into principal components (PCs; Table S5). PC1 and PC2 together described 78% of observed variation and were used here to define climate space independent of location. PC1 was interpreted as ‘continentality’ whereas PC2 reflected a ‘temperature – wetness’ gradient (Fig. S3). Spatial correlations between our climate and demographic niche models were explored by assembling three bins of equal width wherein climate was similar, ensuring the smallest bin had ≥ 100,000 pixels with predictions (Fig. S4). Agreement between maps was estimated as described above.

## Results

### Demographic models

We predicted the growth rate (λ) of song sparrow populations in western North America using a deterministic model based on 48 years of data of juvenile and adult survival (S_j_, S_a_) and reproductive success (RS) and relationships between climatic conditions in our focal population. Eight climate variables hypothesized to influence demography in our focal population accounted for 47, 44 and 56% of variation in annual S_j_, S_a_, and RS observed, respectively (Table 1, S1, S3). S_j_ was predicted to increase with rainfall in August (β ± SE; Aug PPT = 0.254 ± 0.105) and slightly with freezing temperature in December (Dec DD < 0 = 0.190 ± 0.109), but to decline with increasing heat in July (Jul DD > 18 = - 0.390 ± 0.125; (Jul DD > 18)^2^ = 0.195 ± 0.133) and freezing temperatures February (Feb DD < 0 = 0.057 ± 0.137; [Feb DD < 0]^2^ = -0.457 ± 0.106). S_a_ was similarly predicted to increase with rainfall in August (Aug PPT = 0.134 ± 0.059) and freezing temperatures in December, to but decline in with freezing temperatures in February (Dec DD < 0 = 0.112 ± 0.061; Feb DD < 0 = -0.065 ± 0.086; (Feb DD < 0)^2^ = - 0.331 ± 0.082). RS was predicted to increase with increasing warmth and precipitation in spring (Jun DD > 18 = 0.207 ± 0.059; Mar PPT = 0.151 ± 0.055; Jun PPT = 0.175 ± 0.059), factors also influencing the start and duration of breeding.

### Climate niche models

Predicted resident and migrant climate niche models displayed high accuracy (OOB error: 15.5% and 13.9%), sensitivity (81.1% and 83.0%), and specificity (78.4% and 80.3%), respectively (Table S4). Models to predict the resident and migrant climate niches of song sparrows in western North America were trained using 10 climate variables and evaluated by model performance and classification error. Variables reflecting ambient and extreme temperatures had the largest influence on model accuracy in our resident niche model (e.g., DD > 5, Tmax, Tmin, Tave) with DMA ranging from 11.5 to 12.7% (Fig. S2). The predicted migrant niche also depended strongly on local climate, especially temperature in late winter to early spring (Fig. S2; Tmax, DD > 5, Tmin, DD > 18; DMA = 12.4 to 13.5%.) Precipitation also influenced resident and migrant niches strongly (e.g., see below; Fig. S2; DMA = 10.6 and 12.2%, respectively).

However, variables used to develop climate niche models affected migrant and resident niche differently over our study area in western North America (Fig S5). In all cases, warmer winter temperatures prevailed in the resident versus migrant niche, but these differences were much smaller in California than Alaska, causing a statistical interaction between location and migratory status (Table S6). In contrast, winter precipitation was higher in the migrant than resident niche in AK and BC whereas an opposite pattern prevailed in niche CA (Table S6). Variables reflecting heat load in spring also influenced resident and migrant climate niches unequally. For example, degree-days of warming (DD > 18) in the migrant niche in CA exceeded values recorded in the resident niche. Warming was also higher in the resident than migrant niche in BC but similar in migrant and resident niches in AK (Table S6).

### Spatial agreement of demographic and climate niches

The parallel effects of climate on the demography of our focal population and modeled distributions of resident and migratory populations in western North America led to strong agreement of the predicted niches in Cartesian space between our demographic and climate niche models in the region of our focal population (BC; 0.728; Fig. 1c, f). However, agreement among predictions was slightly to much weaker at the northern (AK; 0.627) and southern (CA; 0.259) extents of the study area, respectively. Spatial correlations between the predicted distributions of resident and migratory sparrows also declined as the Euclidean distance from our focal population increased (*r* = -0.24 ± 0.02), as expected if locally adapted populations respond to climatic conditions differently than those resident year-round in our focal study population.

**Figure 1.**
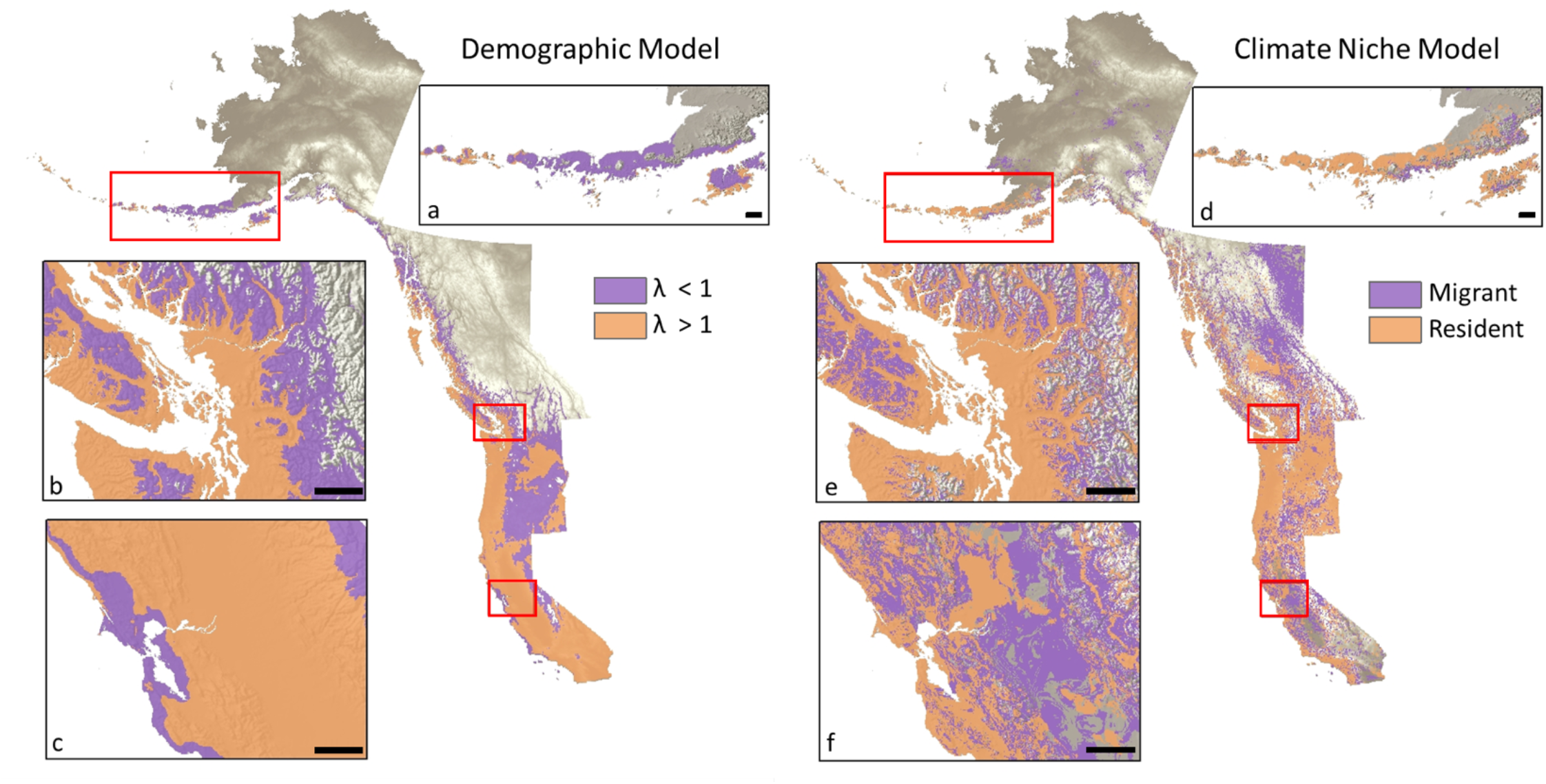
Present-day ranges of the expected population growth wherein the migrant demographic niche is indicated in purple (λ < 1), and resident demographic niche is indicated in orange (λ > 1) predicted using demographic models (insets a-c), and migratory (purple) and resident song sparrow populations (orange) predicted using climate niche models (insets d-f; see Methods). AK (inset a and d; Aleutian Islands, AK), BC (inset b and e; Georgia Basin, BC), and CA (inset c and f; San Francisco Bay, CA) regions highlight areas wherein the observed ranges are more and less similar. Climate data was acquired from ClimateNA (version 6.00; Wang et al. 2016). Scale bar: 50km.

Agreement of our climate and demographic niche models also declined as climatic conditions in our focal population diverged from those experienced by song sparrows elsewhere in western North America. Across western North American variation in climate space was largely accounted for by PC1 (56.2%; Table S5), reflecting differences in ‘continentality;’ e.g., coastal areas with positive values of PC1 experience much less variation in annual temperature and precipitation than montane, high plains, and deserts, all with negative values of PC1 (Fig. S3). PC2 described a complimentary but less variable (21.8%; Table S5) gradient of warmer, drier (positive values) to cooler, wetter conditions (negative values; Fig. S3).

Strong spatial correlations between the demography and predicted distribution of resident and migratory song sparrows at regional to larger scales similar to those estimated between continentality (PC1, Fig S4) and the spatial overlap of the predictions of demographic and climate niche models in the region surrounding our focal population (bin 2: *r* = 0.78 ± 0.01). However, as predicted, we observed smaller positive correlations in the northern region of our study area (AK), where variation in continentality was very high (bin 1: *r* = 0.53 ± 0.02), and in the southern region of our study area (CA) where variation in continentality was less but variation in aridity was high (bin 3: *r* = 0.32 ± 0.02). Despite accounting for a substantial fraction of variation in climate over the study area, PC2 was weakly related to the predicted ranges of resident and migrant song sparrows based on our climate niche model (Fig S4).

### Historical and future variation in the spatial distribution of the climate niche

General agreement of our demographic and climate-based niche models allowed us to ask how migratory and resident niches may have shifted due to climate warming after the period 1901-1910, when winter temperatures at our focal population averaged 0.9°C less than conditions in 2010-2018, but still 3.1°C less than averages expected in 2070-2100. To predict how such changes could affect the demography and distribution of song sparrows in the region of our focal population, we therefore compared our climate and demographic niche models parameterized to historical, contemporary, and future conditions to quantify consequence shifts in the range of resident and migrant song sparrow populations. In particular, we expected that climate warming has facilitated range expansion by song sparrows, particularly those displaying resident life histories.

In support of our predictions, the combined area of the resident and migrant song sparrow niches in the BC region was predicted to have increased by 27.7% (39,668 km^2^; Fig. 2a, b) and 24.1% (36,309 km^2^; (Fig. 2d, e) from 1901-1910 to present (demographic and climate niche model, respectively). The demographic and climate niche model predicted further increases of 27.4% (50,003 km^2^; Fig. 2b, c) and 10.1% (18,511 km^2^; Fig. 2e, f), respectively, given climate in 2070-2100. Similarly, the resident niche was predicted to have increased by 36.3% (24,467 km^2^; Fig. 2a, b) from the historical to present period under our demographic model, or by 38.8% (37,471 km^2^; Fig. 2d, e) in climate niche model. The resident niche was also projected to increase by 12.8% (17,015 km^2^; Fig. 2e, f) from the present to future period by our climate niche model, but by 69.1% under our demographic niche model (63,384 km^2^; Fig. 2b, c). Demographic and climate models also predicted upward expansions in elevation range for song sparrows, with the mean elevation of resident populations rising 171 m over 200 years (i.e., 1900 to 2100; historical = 365 m, current = 475 m, future = 536 m) according to our climate model versus 258 meters (historical = 240 m, current = 281 m, future = 498 m) by our demographic model, confirming close correspondence of these predictive models proximal to our focal study population.

**Figure 2.**
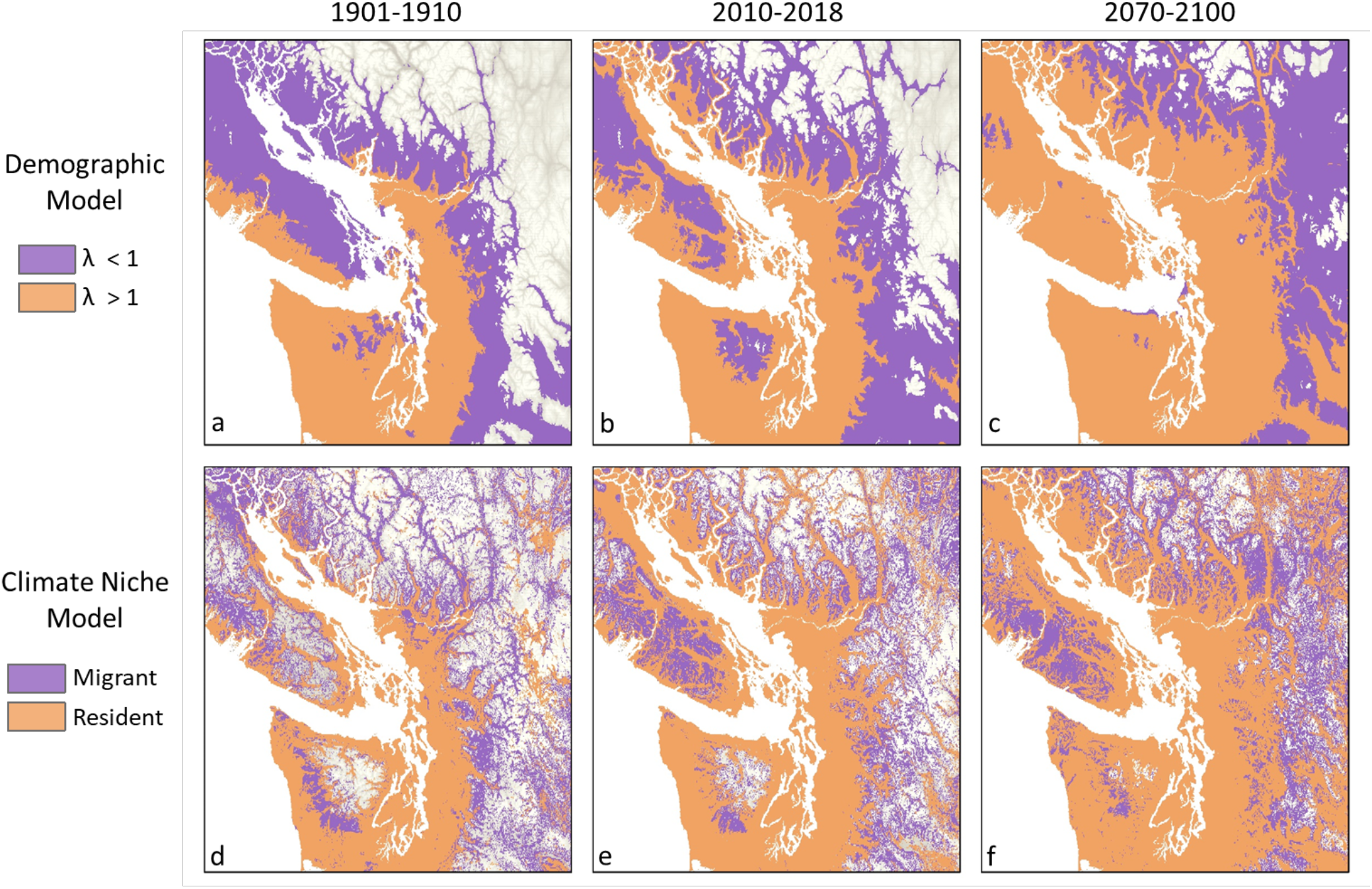
Projected distribution of song sparrows for past (1901-1910; a, d), current (2010-2018; b, e) and future (2070-2100 under scenario RCP 4.5; c, f) using two modelling approaches: demographic modelling (GLM; a-c), and climate niche modelling (Random Forest; d-f) for the BC region. In the demographic models, the migrant niche (λ < 1) is indicated in purple and resident niche (λ > 1) in orange, and in the climate niche models the migrant niche is shown in purple and resident niche in orange. Climate data was acquired from ClimateNA (version 6.00; Wang et al. 2016).

## Discussion

Predicting how species distribution may shift in response to climate change will depend in part on how life-history traits such as seasonal migration versus residence influence species occupancy in response to seasonality and resource availability via their influence on individual fitness and population demography (e.g., Reid et al. 2018, Visty et al. 2018, Acker et al., 2021a; 2021b). Accordingly, we addressed uncertainties in the development and application of species distribution models, their empirical basis, and the reliability of their predictions whilst also exploring the potential for variation in migration to facilitate climate adaptation in a highly polytypic, mobile species. We found substantial agreement in the predictions of demographic and climate niche models in the region encompassing of our focal study population (BC), but lower agreement in areas experiencing different climatic conditions. Our results thus support the idea that climate niche models based on species occupancy and long-term climate averages can reproduce the predictions of detailed, empirical models that link demographic performance to species distribution and migration via space for time substitution. In contrast, our efforts to extrapolate the predicted effects of climate on the demography of our focal study population to song sparrows over their range in western North America revealed mismatches that indicate that the climatic factors driving demography in our focal population differ from those driving the demography of populations in CA and AK. However, our results also suggest that climate has and will continue play a key role affecting the distribution of song sparrows via its influence on demography, and that variation in migration may facilitate climate adaptation by influencing the composition of populations with respect to migratory phenotype and population growth. Taken together, these findings highlight the potential role of life history evolution in climate adaptation while also pointing to limits in our ability to predict long-term change in species distributions and mechanisms involved. We now develop these results with respect to our initial goals.

### Demographic versus climate niches

We found strong agreement between the mapped predictions of empirical models designed to represent the contemporary distributions of resident and migrant song sparrow populations, given assumptions about their response to spatial and temporal variation in climate and its influence on population growth (Fig. 1). These results strongly support the general hypothesis that climate plays a key role in the demography, distribution, and life history of species (Hutchinson, 1957). More importantly, they extend understanding by demonstrating that empirical demographic models driven by climatic conditions can predict migratory phenotype given historical, current, or future climatic conditions (Fig. 2).

Dybala et al. (2013) suggested that the climate niche of song sparrows at a long-term monitoring site in coastal California was strongly influenced by winter temperature and precipitation in the preceding wet season via their effects on survival and population growth. Our demographic model predicted a declining population growth rate (λ= 0.80) in coastal California, similar to Dybala et al.’s (2013) prediction given averaged climatic conditions (λ = 0.88). However, because we arbitrarily classified populations predicted to decline (λ < 1) based on local climate and modeled relationships between demography and climate in our focal population, our demographic niche model predicted that this coastal California population should migrate, whereas our climate niche model accurately predicted their status as year-round residents. It is plausible that some song sparrow populations in coastal California exist as metapopulations comprising a mix of migratory phenotypes. Although, these comparisons suggest our empirical demographic niche model under-predicted the range of resident song sparrows in western North America, they generally support suggestions that climate niche models can be used to elucidate factors contributing to variation in the historical, contemporary, and future distributions of partial migrant species.

Across western North America our climate and demographic niche models yielded moderate to excellent predictions of the migratory status of song sparrow populations (Fig. 1; Table S3; S4). In particular, models built on an *a priori* understanding of the effects of climate on local survival, reproduction, and population growth predicted the observed distribution of resident and migratory populations in the BC region with high precision. This supports the hypothesis that variation in the occurrence of seasonal migration represents an adaptive response to temporal variation in climate and its influence on resource availability and osmotic and/or thermal stress. Moreover, the close correspondence of our empirical climate niche model with observed patterns of spatial variation seasonal migration in BC imply that this variation is shaped by natural selection (e.g., Arcese et al., 2002; Peters et al., 2017; Reid et al., 2018; Delmore et al., 2020).

### Spatio-temporal shifts in resident and migrant niches

Links between climate, demography, and distribution in BC allowed us to also predict that climate warming in the last century should have facilitated range expansion by song sparrows expressing resident phenotypes as the amelioration of winter cold and relaxed its effects on juvenile and adult survival (e.g, Arcese et al. 1992). Similar examples of range expansion and upward shifts in elevation in response to climate warning have been reported in many taxa (birds, La Sorte & Thompson, 2007; insects, Hickling et al., 2005; plants, Holzinger et al., 2008; mammals, Moritz et al., 2008). Although song sparrows have resided year-round in our focal study population since at least 1960 (Tompa, 1963), our demographic niche model suggests that a population comprised of residents would decline rapidly given historical climatic conditions due to a much higher frequency of freezing temperatures in winter. We therefore expect migratory phenotypes to have predominated in our focal population in the early 1900s, as is currently the case in song sparrow populations breeding above 500 m elevation in BC (Fig. 2, our unpubl. observations). Although historical data from BC do not allow a direct test of this hypothesis, a shift from migratory to residential phenotype has occurred at Interpont, Ohio, where Nice (1933) used color-banded birds to confirm the migratory status of all but a single male song sparrow in the 1930s, but wherein most or all birds now reside year-round (Chris Tonra, pers. comm.). Historical records in British Columbia also suggest resident populations of song sparrows were restricted to warm coastal sites on Vancouver Island prior to the 1890s (e.g., Fannin, 1891).

Overall, the distribution of resident song sparrow populations predicted by our models has expanded substantially since historical surveys (Fig. 1), implying a marked expansion over the last century. Similarly, Visty et al. (2018) reported that fox sparrows (*Passerella unalaschcensis*), formerly an obligate migrant throughout its range, established year-round residence on Mandarte Is. after 1960-63, when Tompa (1963) initiated our song sparrow study. Interestingly, Visty et al. (2018) reported that resident fox sparrows produced more broods, and displayed higher survival and population growth rates on Mandarte Is. than migrant populations studied to date, suggesting that higher relative fitness in birds displaying resident versus migrant phenotypes facilitated an expansion of the breeding range.

### Local adaptation

In European shags (*Phalacrocorax aristotelis*), harsh climatic events in winter contributed to mortality and influenced natural selection on the migratory phenotype of individuals, supporting the idea that micro-evolutionary processes can influence the composition of populations with respect to migratory phenotype and affect the breeding distribution of species (Acker et al., 2021a; 2021b). Our results underscore this idea by demonstrating thematic links between climatic, population demography, species distribution, and the evolution of seasonal migration in response to historic and ongoing climate change (Aitken et al., 2008; Sexton et al., 2009; Hendry et al., 2018; Bay et al., 2018). Specifically, we suggest that variation in migration behavior represents one route by which mobile species can adapt to rapid climate change, particularly in partial migrants subject to climatic limits on survival and/or reproduction.

Song sparrows are among the most polytypic vertebrates known (Aldrich, 1984; Arcese et al., 2002; Patten & Pruett, 2009) and vary markedly in migration behavior and correlated behavioral, morphological and physiological traits including body dimensions (Pruett & Winker, 2010), clutch size (Johnston, 1954), and osmoregulatory capacity (Mikles et al., 2020). Song sparrows also vary predictably in traits widely recognized as adaptations to climatic variation in seasonality and primary production (Sæther et al., 2016) including migratory, territorial, dispersal, and breeding behaviors, and demographic traits linked to fecundity, parental effort, and longevity (Arcese, 1989; Arcese et al., 2002; Germain & Arcese, 2014; Tarwater & Arcese, 2017; Reid & Arcese, 2020). Because many such traits have an additive genetic basis (e.g., Schluter & Smith, 1986; Wolak & Reid, 2016; Reid & Arcese, 2020), it is plausible that spatial variation in natural selection has contributed to heritable variation in migratory phenotype, as extensively described in European blackcaps (*Sylvia atricapilla;* e.g., Berthold, 1991; Berthold & Pulido, 1994; Delmore et al., 2020). If so, the pace of adaptation to climate warming in song sparrows might first be measured as the rate by which residency has become established in local populations known to have been migrants historically, and secondarily by estimating the rate of change in allele frequencies at functional loci (Rellstab et al., 2016; Capblancq et al., 2020). Phenotypic changes in migration may also arise plastically and influence subsequent genetic change (Coppack & Pulido, 2004; Pulido, 2007; Teplitsky & Millien, 2014). Quantitative genetic approaches (Kruuk, 2004; Wolak et al., 2018; Reid et al., 2021) to estimate the genetic basis of partial migration and pedigree-based comparison of individuals displaying different migratory tactics should also help elucidate the contributions of phenotypic plasticity versus micro-evolution in local adaptation to rapid environmental change.

### Conclusions

The ability of song sparrows to persist over a wide range of climatic conditions in western North America indicates a broad tolerance for a diverse range of factors with the potential to limit individual fitness and population growth. By engaging in seasonal migration from sites where residents are unlikely to persist year-round, song sparrows encompass a larger range in North America than could otherwise be accommodated. The rapid evolution of populations comprised of migrant, partial migrant, or resident song sparrows, as evidenced by transitions to residency in areas subject to climate warming, thus appears to be a common mechanism by which song sparrows and other mobile species are adapting to local and global climate change. However, regardless of taxa, phenotypic change in migratory status is likely to be accompanied by changes in a suite of physiological, morphological, and behavioral traits influenced by plastic and genetic pathways. Consequently, predicting the speed at which populations can respond to change still requires a better understanding of the loci involved, the influence of natural selection on standing variation in focal and correlated traits, and the genetic structure of such populations. Addressing such uncertainties could advance conservation planning, but is challenging in the absence of robust, long-term studies capable of elucidating the mechanisms underlying local adaptation to climate.

## Acknowledgements

We are deeply grateful to the Tswaout and Tseycum First Nations Bands for allowing us to work on Mandarte Island and to the Natural Sciences and Engineering Research Council of Canada, University of British Columbia, FRBC Chair in Conservation, Hesse Graduate fellowship, and Norwegian Research Council and NTNU Centre for Biodiversity Dynamics (SFF-III 223257) for recent and long-term support.

