## Supplemental Data for "Seasonal migration as a life history trait facilitating adaptation to climate change"

**Supplementary Information**

***Table S1*.** Climate variables used to build demographic niche model for the focal song sparrow population. Abbreviations of the variable names and the range of values for each monthly climate variables as produced by ClimateNA for 2010-2018 are provided below.

| **Variable** | **Abbreviation** | **Min** | **Max** |
| --- | --- | --- | --- |
| August Precipitation (mm) | Aug PPT | 0 | 178.8 |
| February Degree Days < 0°C | Feb DD < 0 | 0 | 94.8 |
| December Degree Days < 0°C | Dec DD < 0 | 0 | 104.6 |
| July Degree Days > 18 °C | Jul DD > 18 | 0 | 81 |
| March Precipitation (mm) | Mar PPT | 0 | 142.9 |
| June Precipitation (mm) | Jun PPT | 0 | 83 |
| June Degree Days > 18 °C | Jun DD > 18 | 0 | 67.3 |

***Table S2*.** Summary of climate variables used to model climate niche of song sparrows in the study area. Abbreviations of the variable names and the range of values for each variable during winter and breeding season as produced by ClimateNA for 2010-2018 are provided below.

|  |  | **Resident** | | **Migratory** | |
| --- | --- | --- | --- | --- | --- |
| **Variable** | **Abbreviation** | **Min** | **Max** | **Min** | **Max** |
| Mean Temperature (°C) | Tave | -27.2 | 16.1 | -18.1 | 32.8 |
| Max. Temperature (°C) | Tmax | -22.9 | 23.8 | -16.2 | 41.3 |
| Min. Temperature (°C) | Tmin | -31.4 | 12.0 | -20.5 | 24.4 |
| Precipitation (mm) | PPT | 0 | 1198.2 | 0 | 787.2 |
| Degree Days < 0°C | DD < 0 | 0.8 | 803.3 | 0 | 383.2 |
| Degree Days > 5°C | DD > 5 | 0 | 325.9 | 0 | 841.2 |
| Degree Days > 18 °C | DD > 18 | 0 | 30.7 | 0 | 440.7 |
| Number of Frost-Free Days | NFFD | 0 | 29.5 | 0 | 30.5 |
| Relative Humidity | RH | 38.9 | 98.8 | 36.6 | 100.0 |
| Precipitation as Snow (mm) | PAS | 0 | 1178.9 | 0 | 601.3 |

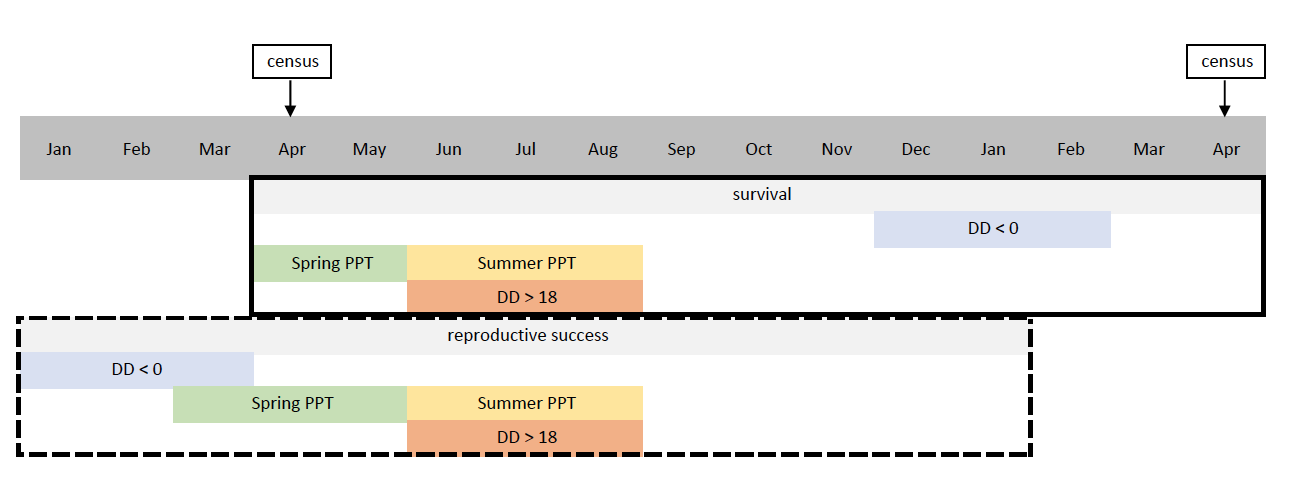

***Figure S1.*** A timeline showing when variables used in the demographic models were measured. A census of the population took place every April in which all birds on the island were counted. Adult survival was measured between censuses and juvenile survival was measured between the time of fledging and the next census. Colored variables represent climate variables hypothesized to influence vital rates.

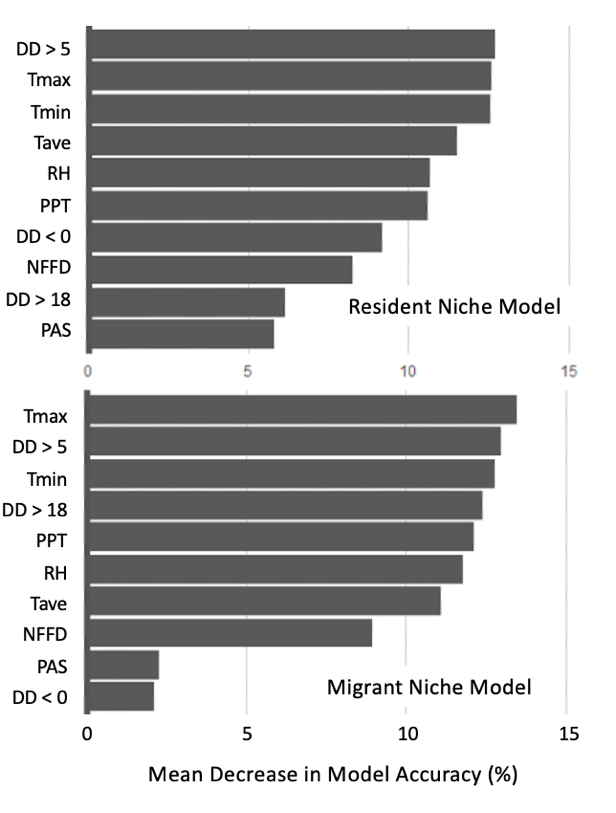

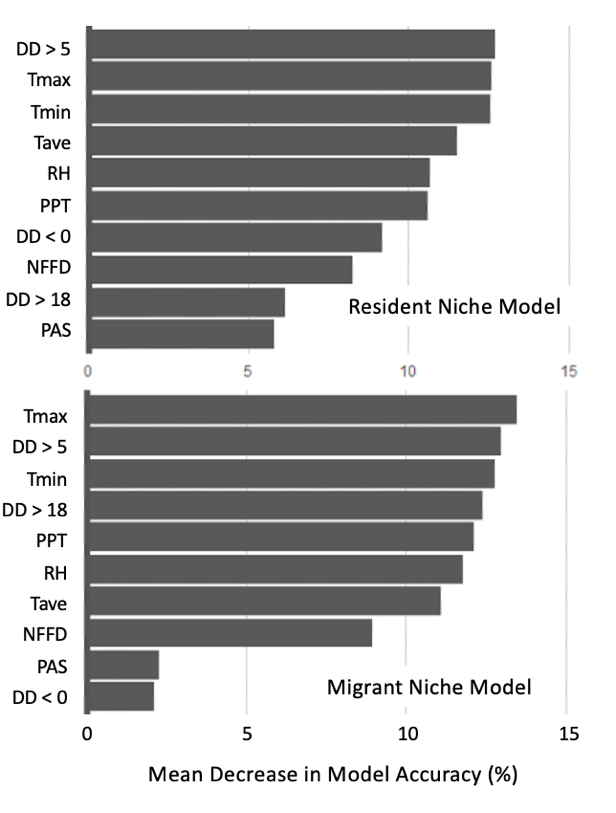

***Fig S2*.** The variable importance of climate used in the climate niche models. Variables are ordered by ranked importance, as the mean decrease in model accuracy (DMA) in percent.

| ***Table S3.*** Formulas and GLM model output for vital rate models used to calculate the demographic niche model. | | | | | | | |
| --- | --- | --- | --- | --- | --- | --- | --- |
| **Vital Rate** | **GLM Formula** | **Output** | **RMSE** | **Variable** | **Estimate** | **SE** | **Variable Importance** |
| Adult Survival | Aug PPT + Feb DD < 0 + (Feb DD < 0)^2^ + Dec DD < 0 | F(4,41) =12.89,  p < 0.001,  R^2^ = 0.56 | 0.359 | Aug PPT | 0.134 | 0.059 | 2.281 |
|  |  |  |  | Feb DD < 0 | -0.065 | 0.086 | 2.201 |
|  |  |  |  | (Feb DD < 0)^2^ | -0.331 | 0.082 | 3.883 |
|  |  |  |  | Dec DD < 0 | 0.112 | 0.061 | 1.825 |
| Juvenile Survival | Aug PPT + Feb DD < 0 + (Feb DD < 0)^2^ + Dec ^o^ Days < 0 + Jul DD > 18 + (Jul DD > 18)^2^ | F(6,39) =5.14,  p < 0.001,  R^2^ = 0.44 | 0.598 | Aug PPT | 0.254 | 0.105 | 2.413 |
|  |  |  |  | Feb DD < 0 | 0.057 | 0.137 | 1.757 |
|  |  |  |  | (Feb DD < 0)^2^ | -0.457 | 0.106 | 2.568 |
|  |  |  |  | Dec DD < 0 | 0.190 | 0.109 | 1.735 |
|  |  |  |  | Jul DD > 18 | -0.390 | 0.125 | 1.699 |
|  |  |  |  | (Jul DD > 18)^2^ | 0.195 | 0.133 | 1.097 |
| Reproductive Success | Mar PPT + Jun PPT + Jun DD >18 | F(3,43) =8.49,  p < 0.001,  R^2^ = 0.47 | 0.365 | Mar PPT | 0.151 | 0.055 | 2.749 |
|  |  |  |  | Jun PPT | 0.175 | 0.059 | 2.929 |
|  |  |  |  | Jun DD > 18 | 0.207 | 0.059 | 3.498 |

***Table S4.*** Predictive performance metrics used to compare the predictions to the actual observations from unseen data (out-of-bag samples): OOB error, sensitivity, specificity, AUC, and Kappa for the climatic niches of resident and migrant song sparrows.

| **Model** | **OOB error** | **Sensitivity** | **Specificity** | **AUC** | **Kappa** |
| --- | --- | --- | --- | --- | --- |
| Resident niche model | 0.155 | 0.811 | 0.784 | 0.886 | 0.591 |
| Migrant niche model | 0.139 | 0.830 | 0.803 | 0.896 | 0.626 |

***Table S5.*** Climatic variables used to define climate space of the study area and loadings of each variable on the first two principal component axes that explained 78% of the variation (PC1: 56.2%; PC2: 21.8%). Monthly data from ClimateNA for the contemporary period (2010-2018) during the winter season (Jan – Feb) were aggregated into averages. See Fig. S2 for maps of the loading of PC1 and PC2.

| **Variable** | **Loadings** | |
| --- | --- | --- |
|  | PC1 | PC2 |
| Tmax | 0.407 | 0.049 |
| Tmin | 0.412 | -0.054 |
| Tave | 0.412 | -0.002 |
| PPT | 0.224 | -0.486 |
| DD_0 | -0.389 | 0.11 |
| DD5 | 0.293 | 0.399 |
| DD18 | 0.193 | 0.33 |
| NFFD | 0.373 | 0.182 |
| PAS | 0.044 | -0.519 |
| RH | 0.17 | -0.419 |

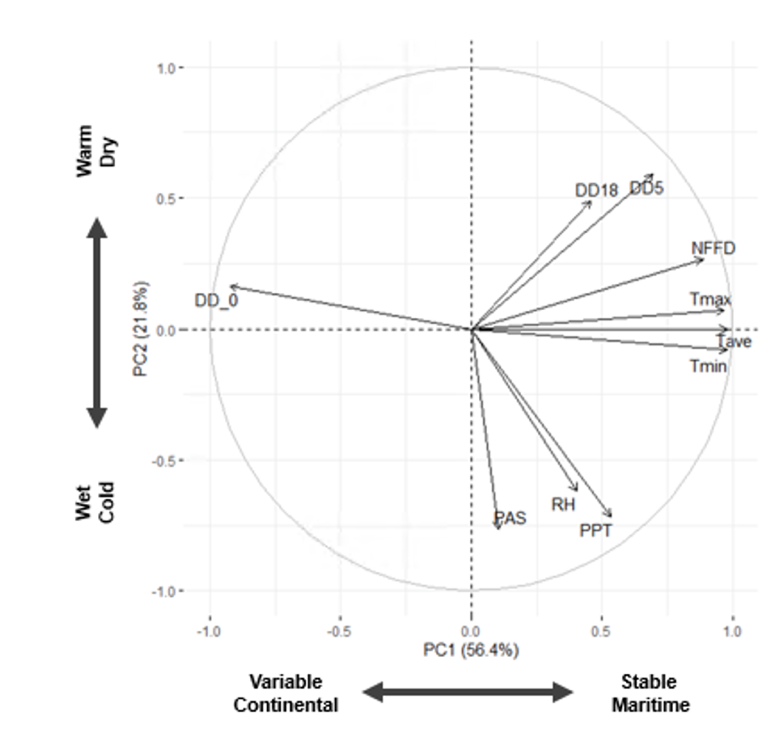

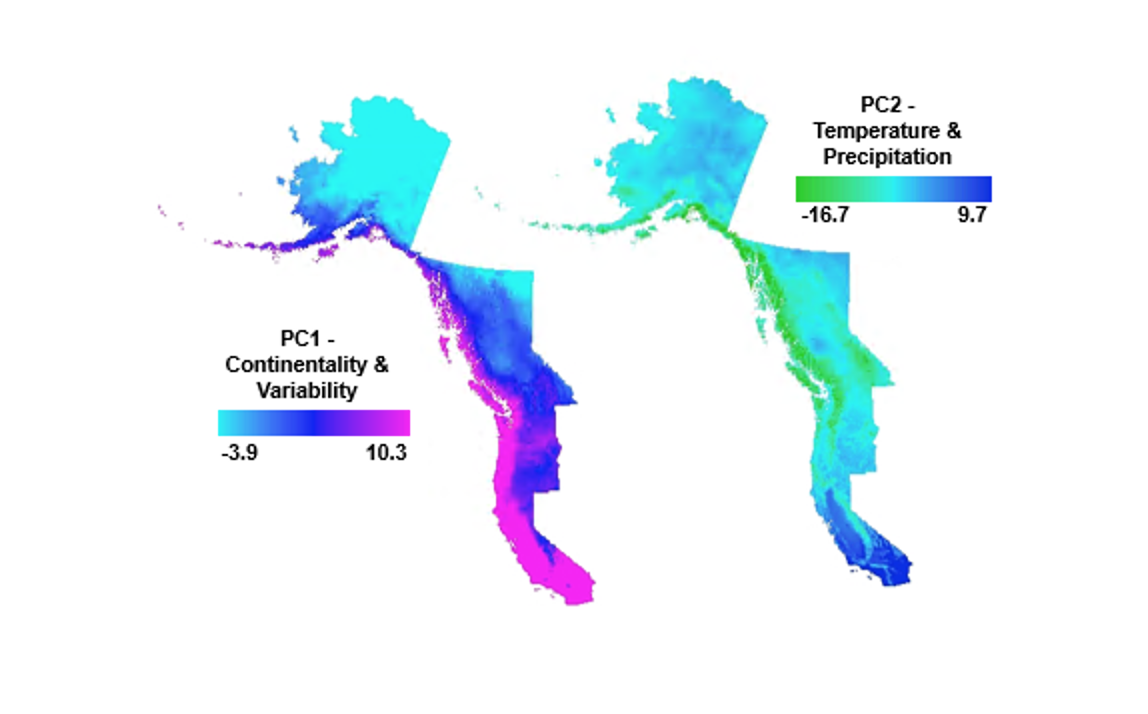

**B**

**C**

**A**

***Figure S3.*** The primary climate gradients in the study area identified through a principal component analysis of the contemporary climate variables during winter. (A) Biplot of the climate gradients of variability and continentality, and temperature and precipitation, as represented by components (PC) 1 and 2. Spatial distribution of the variability and continentality gradient scores (B; PC1), and temperature and precipitation gradient scores (C; PC2). Variables were standardized prior to the PCA so that each one has mean zero and unit variance independent of its scale to ensures that all variables have the same weight in the analysis. Abbreviations of the climate variables shown in the biplot (A) are explained in Table S2.

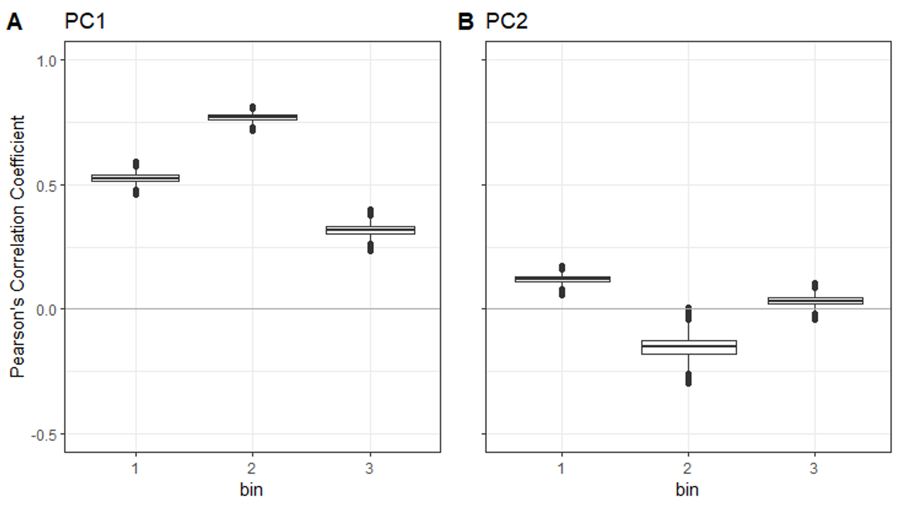

***Figure S4.*** The estimated Pearson’s correlation (*r*) of predictions from the contemporary resident climate niche model and demographic model of the study region grouped in bins where climate is considered similar. PC1 (A) and PC2 (B) represents variability and continentality gradient scores and temperature and precipitation gradient scores, respectively.

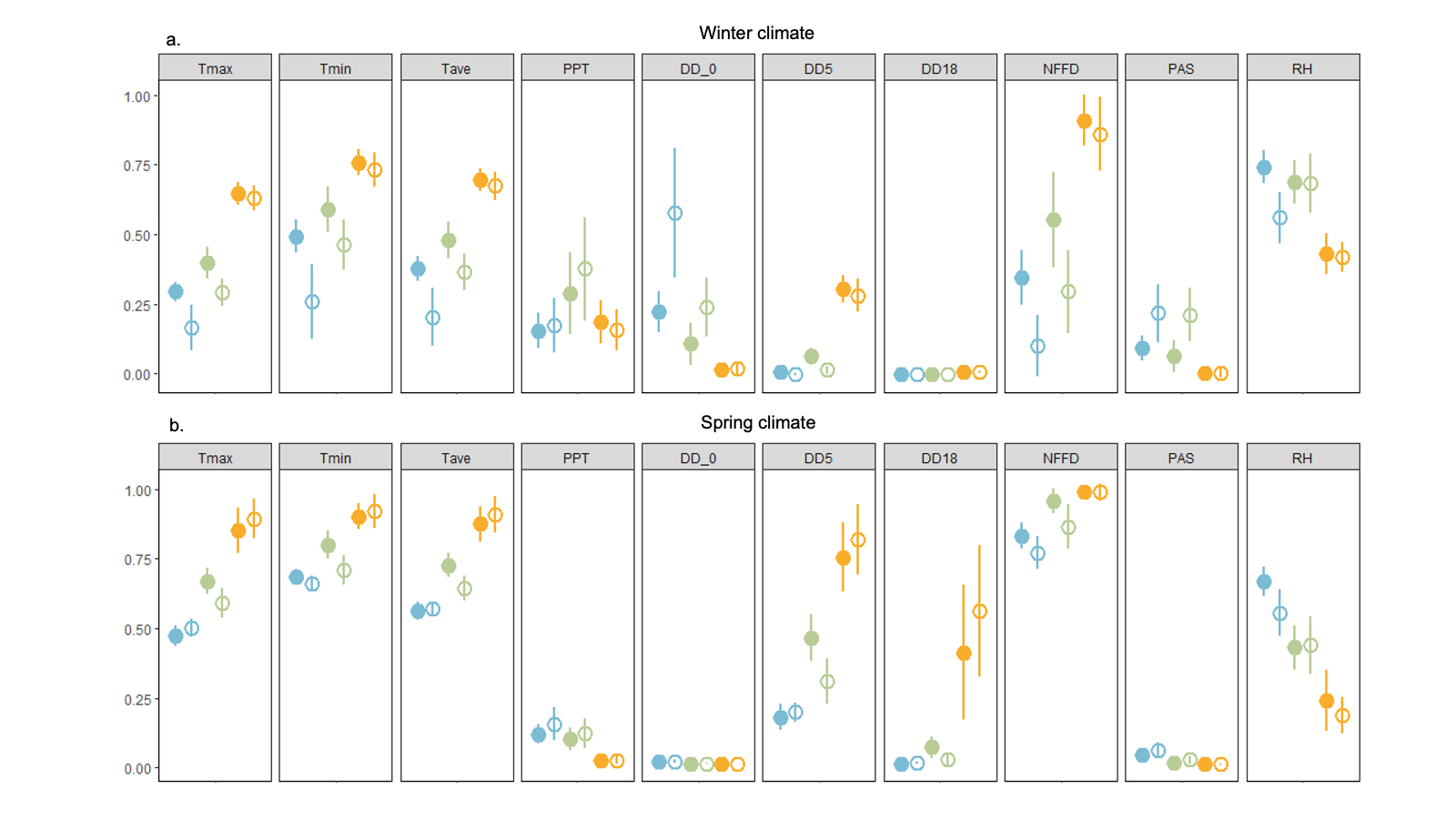
***Figure S5.*** Variation (mean ± SD) in the variables used in the Climate Niche Model by inset locations shown in figure 1 in winter (a) and spring (b): Aleutian Islands, AK (blue), Georgia Basin, BC (green), and San Francisco Bay, CA (yellow). Filled and open circles represent the resident and migratory niches, respectively. To facilitate visual comparison variables were normalized to bring values to range from 0-1 by subtracting the minimum and dividing by the maximum and sampled randomly within each location and each niche (*n* = 5,000 per niche per location).

**Table S6.** F-statistics from the analysis of variance (ANOVA) used to describe the mean and range of variation in the contemporary migrant and resident niche climate variables. Raw climate variables used in the Climate Niche Model were randomly sampled by inset location shown in Figure 1 for each niche in winter and spring (*n* = 5,000 per niche per location). All values reported were significant at p < 0.001.

|  |  | **Tmax** | **Tmin** | **Tave** | **PPT** | **DD < 0** | **DD > 5** | **DD > 18** | **NNFD** | **PAS** | **RH** |
| --- | --- | --- | --- | --- | --- | --- | --- | --- | --- | --- | --- |
| **winter** | niche | 17798.1 | 17377.3 | 18396.4 | 435.4 | 15747.7 | 4523.0 | 534.7 | 15236.1 | 15070.5 | 5175.2 |
|  | location | 151961.0 | 48459.8 | 92755.8 | 6615.1 | 28785.7 | 210594.0 | 49615.7 | 70189.6 | 16228.0 | 32558.6 |
|  | niche:location | 2956.4 | 3807.7 | 3450.5 | 671.9 | 6127.4 | 883.7 | 280.6 | 2106.6 | 3655.2 | 4153.2 |
| **spring** | niche | 6.7 | 3544.6 | 692.2 | 1889.1 | 1044.9 | 587.9 | 477.8 | 8089.6 | 3476.5 | 2870.7 |
|  | location | 120860.0 | 69989.6 | 117301.1 | 22280.1 | 30045.1 | 115815.1 | 35998.3 | 35421.0 | 24387.3 | 55596.4 |
|  | niche:location | 3546.6 | 3730.7 | 4171.3 | 532.9 | 271.3 | 4263.1 | 1291.9 | 2057.2 | 857.7 | 1340.2 |

**Workflow**

1. Response variables (S_a,_ S_j_, RS) were mean-centered and natural-log transformed.
2. The full linear model included all variables in Table 1 (linear and second-order polynomial terms) with the transformed response variable from step 1.
3. We reduced the full model via supervised backward selection until only variables with p < 0.1 were remaining for our reduced models (Table S3).
4. Using *raster* package, we used our reduced models to predict these relationships across our study area. The resulting predictions were then back-transformed as follows:

e ^Sa – 0.6607^

e ^Sj – 1.73617^

e ^RS + 0.4596^

5. We then constrained predicted values between 3 ± SE of mean observed in our focal population on Mandarte, Island as follows:

S_a_: 0 – 0.99

S_j_: 0 – 0.69

RS: 0 – 4.34
